# The *Escherichia coli* small heat shock protein IbpA plays a role in regulating the heat shock response by controlling the translation of σ^32^

**DOI:** 10.1101/2023.03.28.534623

**Authors:** Tsukumi Miwa, Hideki Taguchi

## Abstract

Small heat shock proteins (sHsps) act as ATP-independent chaperones that prevent irreversible aggregate formation by sequestering denatured proteins. IbpA, an *Escherichia coli* sHsp, functions not only as a chaperone but also as a suppressor of its own expression through posttranscriptional regulation, contributing to negative feedback regulation. IbpA also regulates the expression of its paralog, IbpB, in a similar manner, but the extent to which IbpA regulates other protein expressions is unclear. We have discovered that IbpA downregulates the expression of many Hsps by repressing the translation of the heat shock transcription factor σ^32^. The IbpA regulation not only controls the σ^32^ level but also contributes to the shut-off of the heat shock response. These results revealed an unexplored role of IbpA to regulate heat shock response at a translational level, which adds a new layer for tightly controlled and rapid expression of σ^32^ on demand.

**Significance Statement:** To survive during heat shock, cells have a mechanism to induce the synthesis of Hsps and to restore normal levels when the stress subsides. The molecular mechanisms of the heat shock response in *E. coli* have been extensively studied over the years. The master heat shock transcriptional regulator, σ^32^, which is normally at low levels due to chaperone-mediated degradation, is increased upon heat shock. Our study has identified a previously unknown factor, IbpA, that regulates the level of σ^32^ by suppressing its expression at a translational level, thereby contributing to the heat shock response regulation.

## Introduction

Heat stress responses are vital to prevent heat stress-induced protein homeostasis (proteostasis) failure. Coping with stress-induced protein denaturation, Hsps, including chaperones and proteases, act through multiple quality control mechanisms, such as sequestration, refolding, and degradation (1, 2). Hsps are expressed in response to heat stress and are typically regulated at the transcriptional level in both prokaryotes and eukaryotes (1).

Heat stress responses are primarily regulated by changes in the levels or activity of transcriptional regulators (3, 4). In *Escherichia coli*, the heat shock transcription factor σ^32^ regulates the expression of approximately 100 genes (5). Its abundance is immediately increased under heat stress, reaching a maximum at 5 minutes, and gradually decreases thereafter (3, 6, 7). The cells then enter a shut-off phase where σ^32^ is suppressed in both abundance and activity (3, 6, 7). Previous studies on σ^32^ have revealed that its regulation occurs at multiple levels, including transcription, translation, stability, and activity. In particular, posttranscriptional regulations play a significant role in the σ^32^ stress response (3, 6–10).

Chaperone binding to σ^32^ regulates its stability and activity (3, 11, 12). Two major cytoplasmic chaperone systems, DnaK-DnaJ and GroEL-GroES, play various roles, including assisting in the folding of nascent polypeptide chains and maintenance of folded and denatured proteins. Chaperone binding to σ^32^ inhibits its activity and assists in its degradation by the inner membrane protease FtsH (2, 3, 6, 12–14). The membrane localization of σ^32^ via signal-recognition particle (SRP) and its receptor is also essential for the σ^32^ degradation by chaperones and FtsH (15, 16). The σ^32^ recruitment by the SRP receptor is followed by the chaperone-dependent degradation. The σ^32^ I54N mutant, a hyperactive mutant, which disrupts the interaction between σ^32^ and SRP, is not inactivated by chaperones and is resistant to the degradation by FtsH (12, 15–17). The abundance of σ^32^ is usually suppressed to less than 50 molecules in the cell, due to a rapid degradation mediated by chaperones and proteases (7). However, during heat stress, stress-induced chaperone recruitment to denatured proteins allows for a transient increase in the abundance of σ^32^ (7). The subsequent transition to the shut-off phase is thought to depend on the accelerated degradation of σ^32^ mediated by the excess chaperones (18, 19). However, it has been reported that the transition to the shut-off phase is also normal in *E. coli* mutant strains including an *ftsH* deleted strain (20, 21). Additionally, even *E. coli* with the hyperactive σ^32^ I54N mutant exhibits a reduction in the level of heat shock response within 15 minutes of the onset of heat shock (17). These observations suggest that the degradation of σ^32^ alone does not fully explain the shut-off mechanism.

The translation of *rpoH*, which encodes σ^32^, is regulated by thermoresponsive mRNA secondary structures called RNA thermometers (RNATs) (3, 22, 23). At normal or low temperatures, the RNATs adopt complex structures, including a region encompassing the Shine-Dalgarno (SD) sequence in the 5’ untranslated region (5’ UTR), close to the translational initiation sites. However, heat fluctuations cause the melting of the structures that mask the SD sequence, enabling the translation of σ^32^ (23). The RNAT of *rpoH* consists of the 5’ UTR and part of the ORF, and it is known that the RNA secondary structure masking the start codon opens in a temperature-dependent manner (3, 22–24). RNATs have been extensively studied in *rpoH* and small heat shock proteins (sHsps) (12, 23). The RNATs in sHsps exhibit conserved shapes in α- and γ-proteobacteria (23, 25, 26). Recently, the RNA thermometer shape of IbpA, an *E. coli* sHsp, has been demonstrated to be a crucial factor in the negative-feedback regulation of IbpA at the translational level (27).

sHsps are a group of low subunit molecular weight (12-43 kD) ATP-independent chaperones that serve as the first line of defense against protein aggregation stress in organisms of all kingdoms (28–30). sHsps have two chaperone functions: preventing the irreversible aggregation of denatured client proteins by binding to them, and aiding in their folding through other chaperones such as DnaK/DnaJ (28–30). Under normal conditions, sHsps form dynamic oligomers with no fixed number of subunits. The oligomeric states are thought to restrict the client-binding region inside the oligomers and to maintain the sHsps in a storage state. The IX(I/V) motif in the C-terminal region of sHsps is responsible for the formation of oligomers (28–30).

IbpA and IbpB, sHsps of *E. coli*, exhibit a high degree of homology and form hetero-oligomers (31, 32). They function as sHsps by sequestering denatured proteins and facilitating their refolding with the aid of other chaperones. While IbpA has a specialized role in substrate binding to prevent irreversible protein aggregation, its substrate release efficiency for transfer to other chaperones is lower than that of IbpB (31–33).

Recently, we discovered that IbpA serves not only as a chaperone but also as a regulator of gene expression. Specifically, IbpA represses its own translation, as well as that of IbpB, by binding to the corresponding mRNAs (27). Under conditions of increased intracellular protein aggregation, IbpA is recruited for sequestration, which releases the repression of translation. This mechanism represents a negative feedback response that is partly assisted by RNA degradation by RNase (27). An oligomer-forming motif (IX(I/V)) of IbpA is critical for this regulation and for mRNA binding. The RNA secondary structure of the 5’ UTR is also essential for regulation, as mutation of the stem-loop structure in the 5’ UTR of the *ibpA* mRNA abolishes the IbpA-mediated expression regulation (27). Furthermore, IbpA also regulates the expression of IbpB, and the RNAT structure of the 5’ UTR of the *ibpB* mRNA, which is not identical to that of the *ibpA* mRNA (25, 26), suggests the possibility that IbpA may recognize other mRNAs to regulate gene expression.

Here, we searched for novel regulatory targets of IbpA with similar molecular mechanisms. A proteome analysis of *E. coli* cells overexpressing IbpA revealed that the abundance of many Hsps was reduced at the transcriptional level, not the translational level. We, therefore, focused on σ^32^, the master transcriptional regulator of Hsps in *E. coli*. The analysis showed that σ^32^ is a target for the translational regulation of IbpA. We found that the IbpA-mediated inhibition of the *rpoH* translation affected changes in the σ^32^ level during the shut-off phase, although it had no effect on the transient increase in the σ^32^ level upon heat shock. The expression level of IbpA slightly influenced the growth of cells after the recovery from heat stress. Our findings suggest a model in which the σ^32^-mediated shut-off is regulated by IbpA at a translational level. Our study focusing on IbpA has illuminated the involvement of this previously unrecognized factor in the established molecular mechanism of the heat shock response in *E. coli*.

## Results

### Overexpression of IbpA leads to downregulation of multiple heat shock proteins

IbpA regulates the expression of itself and its paralog, *ibpB*, at the post-transcriptional level (27). The stem loops present in the 5’ UTR of the mRNA of *ibpA* or *ibpB* constitute a key regulatory element. While secondary structures with multiple stem-loops are common in the 5’ UTRs of *ibpA* and *ibpB*, their sequences and structures are not identical (25, 26). Therefore, we hypothesized that, apart from self-regulation, IbpA may also influence the expression of other proteins in *E. coli*. To identify potential targets of IbpA-mediated regulation, we conducted a mass spectrometry (MS)-based quantitative proteomics analysis to identify proteins, whose expression is altered in *E. coli* upon IbpA overexpression. We compared the proteomes of cells overexpressing IbpA and those overexpressing GFP as a control and found that 41 and 40 proteins were specifically increased (>1.67-fold) and decreased (<0.6-fold), respectively, upon IbpA overexpression (Fig. 1A, Supplementary Dataset S1). Gene ontology analysis revealed that the most significantly altered and statistically significant category in IbpA-overexpressed cells was the group of proteins associated with chaperones. (red circles in Fig. 1A, Fig. S1A, Supplementary Dataset S2). The proteome of cells overexpressing a non-functional IbpA mutant, IbpA_AEA_, which is defective in both oligomer formation and self-suppression activity, did not show the enrichment of chaperone-related proteins among the decreased proteins (Fig. S1B, C, Supplementary Dataset S2), further suggesting that the reduction of chaperones was specific to functional IbpA.

**Fig 1.**
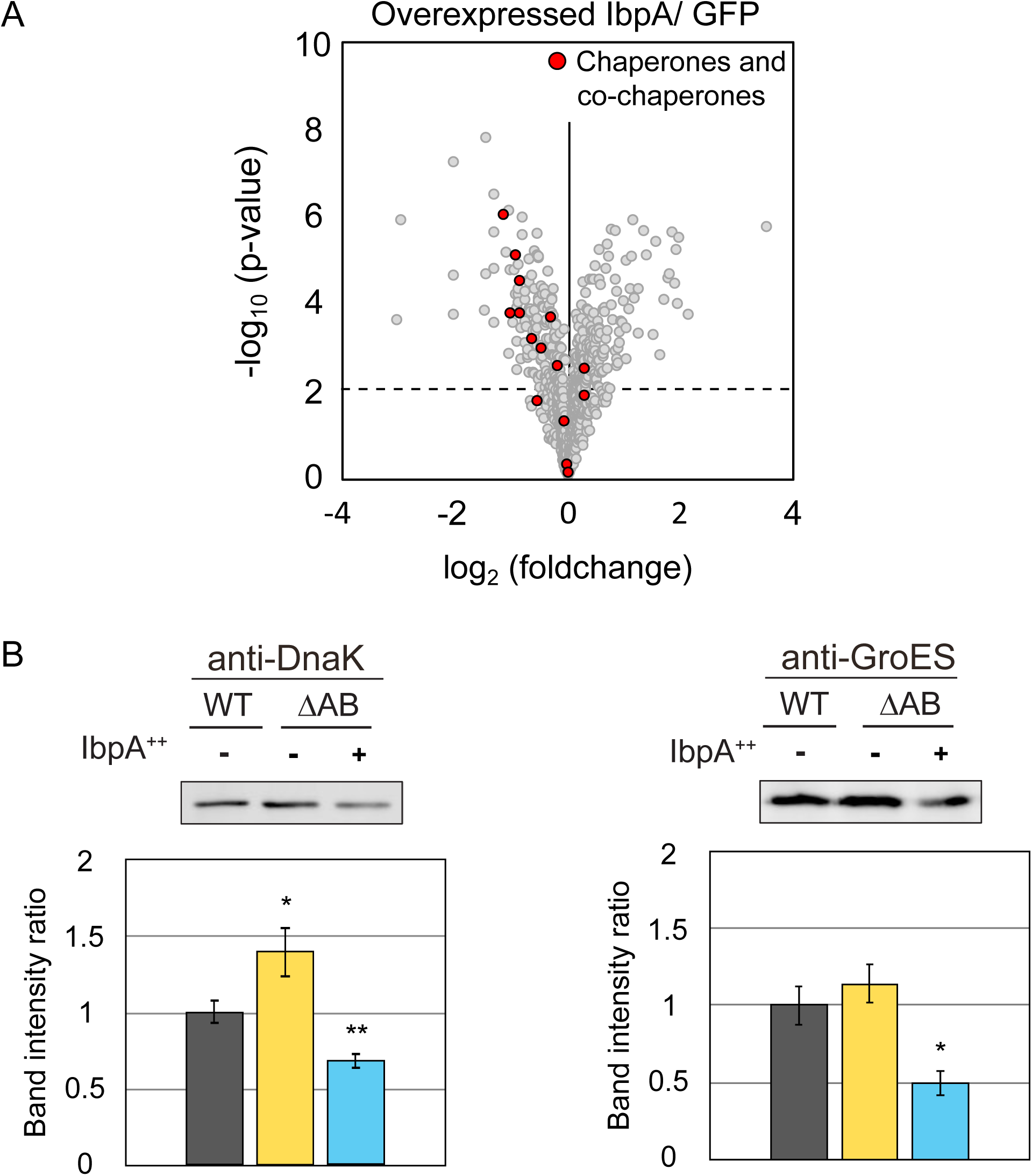
IbpA overexpression downregulates a multitude of heat shock proteins. (*A*) A volcano plot depicting the degree of protein expression ratio between IbpA and GFP overexpression in *E. coli* wild-type strain. Each dot in the plot represents the fold change and *p*-value of an individual protein, identified by comparative proteomics analysis. The dashed line represents a *p*-value threshold of 0.01. Red dots indicate proteins belong to the “chaperone binding” and “unfolded protein binding” categories in gene ontology (GO) term enrichment analysis. (*B*) Western blotting of endogenous DnaK and GroES in *E. coli* wild-type strain (WT) or the *ibpAB* operon-deleted strain (Δ*AB*). IbpA^++^, *E. coli* Δ*AB* cells overexpressing IbpA. Relative band intensities of three biological replicates are shown below. The value in the *E. coli* wild-type was set to 1. Error bars represent the standard deviation (SD); The statistical significance of differences was assessed using the Student’s *t*-test (\**p* < 0.05, \*\**p* < 0.01).

To validate the decrease of chaperones observed in the MS data, we examined the expression levels of representative chaperones, DnaK and GroES by Western blotting. In this experiment, we overexpressed IbpA in an *ibpA-ibpB* deletion strain (Δ*ibpAB*) to better highlight the effect of IbpA abundance; We postulated that depletion of IbpA could lead to an increase in chaperones. The expression level of DnaK and GroES increased in the Δ*ibpAB* strain and decreased in cells overexpressing IbpA (Fig. 1B), supporting our postulation.

## IbpA represses the expression of Hsps at the transcriptional level

Next, to investigate whether IbpA mediates the reduction of chaperones by translation suppression via the 5’ UTRs, we constructed *gfp* reporters harboring 5’ UTRs of *dnaK* and *groES* and examined their expression levels in response to IbpA. We used an arabinose-inducible promoter in the plasmids to keep the mRNA levels almost identical. The abundance of IbpA did not significantly affect the expression of both reporters (*dnaK*: Fig. 2A, *groES*: Fig. S2A). To directly assess the IbpA effect on the *dnaK* and *groES* translation, we used an *E. coli* reconstituted cell-free translation system, PURE system, that only includes factors essential for translation (34). We compared the *dnaK* and *groES* mRNA translation with or without purified IbpA and found that IbpA had no effect on the translation of the *dnaK* and *groES* mRNAs (*dnaK*: Fig. 2B, *groES*: Fig. S2B).

**Fig 2.**
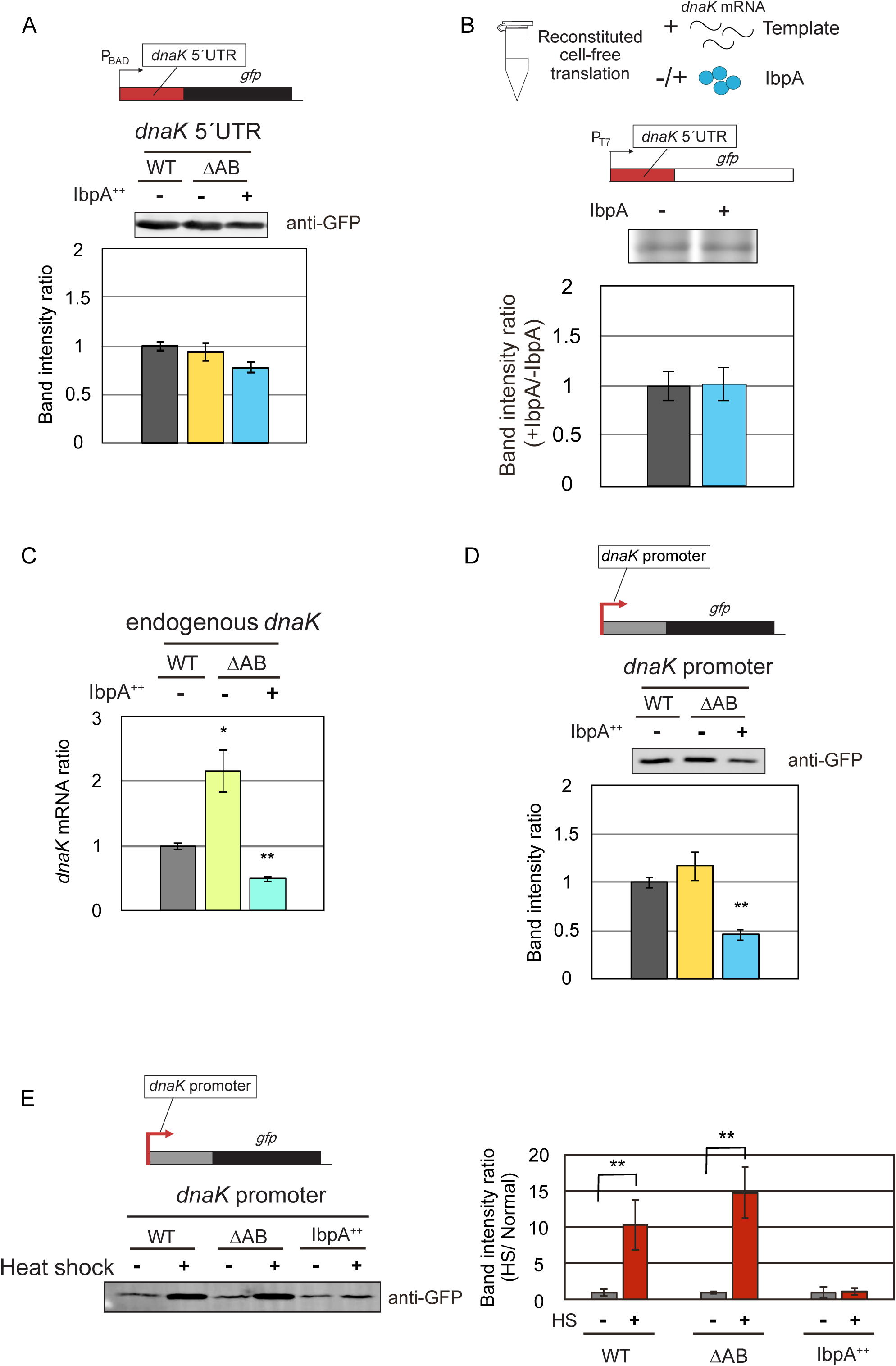
IbpA suppresses the expression of DnaK at a transcriptional level. (*A*) Western blotting of GFP expressed from a *gfp* gene harboring the *dnaK* 5’ UTR in *E. coli* wild-type (WT) or the *ibpAB* operon-deleted strain (Δ*AB*). IbpA^++^, *E. coli* Δ*AB* cells overexpressing IbpA. Relative band intensities of three biological replicates are shown below. The value in the *E. coli* wild-type was set to 1. (*B*) Cell-free translation of the *gfp* reporter in the absence (-) or the presence (+) of purified IbpA using a reconstituted protein synthesis system (PURE system). The value without IbpA was set to 1. (*C*) Quantification of endogenous *dnaK* mRNA levels by qRT-PCR in wild-type cells (*left*), Δ*AB* cells (*middle*), and Δ*AB* cells overexpressing IbpA (*right*). The value in the *E. coli* wild-type was set to 1. (*D*) Western blotting of GFP expressed under the control of the endogenous *dnaK* promoter in *E. coli* cells. Relative band intensities of three biological replicates are shown below. The value in the *E. coli* wild-type was set to 1. (*E*) Western blotting (*left*) and the relative band intensities (*right*) of GFP expressed by the endogenous *dnaK* promoter in *E. coli* cells with (*red*) or without (*gray*) heat shock (*HS*). The values without heat shock were set to 1. Statistical analysis: Error bars represent SD*; n* = 3 biological replicates. The statistical significance of differences was assessed using the Student’s *t*-test (\**p* < 0.05, \*\**p* < 0.01).

We then hypothesized that IbpA may regulate the transcription of *dnaK* and *groES*, and quantified their endogenous mRNA levels in *E. coli* using quantitative real-time PCR (qRT-PCR). The results showed that IbpA overexpression led to a decrease in *dnaK* and *groES* mRNA levels, while the deletion of *ibpAB* increased the levels of these mRNAs (*dnaK*: Fig. 2C, *groES*: Fig. S2C). Furthermore, overexpression of IbpA effectively suppressed the reporter expression in plasmids harboring endogenous *dnaK* or *groES* promoters (*dnaK*: Fig. 2D, *groES*: Fig. S2D). These results suggest that excessive amounts of IbpA repress the expression of DnaK and GroES at the transcriptional level.

The data obtained from the proteome and reporter assay prompted us to hypothesize that IbpA may influence the expression levels of σ^32^, which is the primary transcriptional regulator of Hsps. To investigate this, we examined whether heat stress could impact the expression levels of *dnaK* and *groES* in *E. coli* with varying amounts of IbpA. We observed a substantial increase in the expression of the GFP reporter with the endogenous *dnaK* or *groES* promoter at 42 °C in the Δ*ibpAB* cells, similar to the wild-type. In contrast, such heat stress induction was completely lost in the IbpA-overexpressing cells (Fig. 2E, Fig. S2E). Our findings thus far provide evidence that overexpression of IbpA suppresses the transcription of Hsps by σ^32^. In fact, proteome data revealed that proteins regulated by σ^32^ were overall decreased in cells overexpressing wild-type IbpA, but not in cells overexpressing IbpA_AEA_ (Supplementary Dataset S2), indicating that functional IbpA down-regulates σ^32^ at a translational level.

### Expression of σ^32^ is repressed by IbpA at a translational level

To determine whether the abundance of IbpA can impact the expression level of σ^32^ at a translational level, we examined the endogenous σ^32^ expression in Δ*ibpAB* and IbpA-overexpressing cells. We observed an increase in the expression of endogenous σ^32^ in Δ*ibpAB* cells and a decrease in IbpA-overexpressing cells (Fig. 3A). The drastic reduction in the σ^32^ level was partially due to the decreased *rpoH* mRNA levels in IbpA-overexpressing cells (Fig. S3A), though the degradation rates of the *rpoH* mRNA were consistent in all strains tested (Fig. S3B), indicating that the *rpoH* mRNA level is also influenced by IbpA overexpression. While the *rpoH* mRNA level was not affected in Δ*ibpAB* cells (Fig. S3A), the σ^32^ level was significantly up-regulated in these cells (Fig. 3A). We concluded that the abundance of IbpA affects the σ^32^ level at a translational level. To confirm this, we investigated whether the σ^32^ levels were dependent on the presence of the 5’ UTR of the *rpoH* mRNA using an arabinose promoter to keep the mRNA levels almost identical. The σ^32^ levels were found to be dependent on the 5’ UTR; the trend of the change in the reporter plasmids was the same as that of endogenous σ^32^ levels (Fig. 3B), strongly suggesting that the IbpA suppresses the expression of σ^32^ at a translational level. Finally, in the PURE system, purified IbpA suppressed the translation of the *rpoH* mRNA by approximately 50% in an *rpoH* 5’ UTR-dependent manner (Fig. 3C). In conclusion, our findings demonstrate that IbpA directly suppresses the translation of the *rpoH* mRNA.

**Fig 3.**
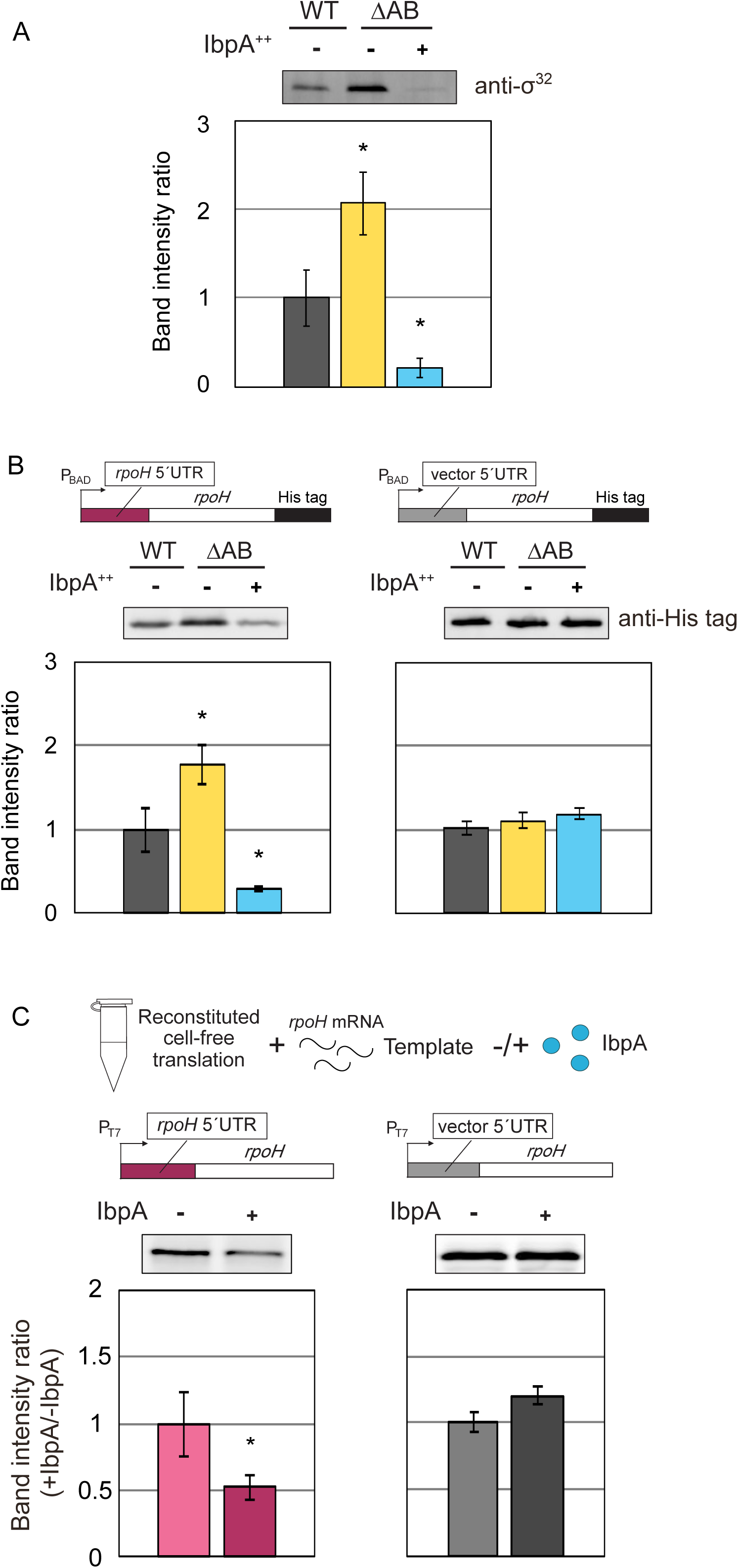
IbpA-mediated suppression of *rpoH* translation. (*A*) Detection of endogenous σ^32^ levels in wild-type and *ibpAB* deleted (Δ*AB*) cells using Western blotting. IbpA^++^, *E. coli* Δ*AB* cells overexpressing IbpA. Anti-σ^32^ antibody was used for the detection. Relative band intensities of three biological replicates are shown below. The value in the *E. coli* wild-type was set to 1. (*B*) Detection of σ^32^ fused with His-tag expressed from plasmids harboring 5’ UTRs of *rpoH* or vector under the control of an arabinose promoter by Western blotting. Anti-His tag antibody was used for the detection. The value in the *E. coli* wild-type was set to 1. (*C*) Cell-free translation of σ^32^ from the *rpoH* mRNA harboring 5’ UTRs of *rpoH* or vector in the absence (-) or the presence (+) of purified IbpA using the PURE system. Cy5-labeled methionines in the translation products were detected with a fluorescence imager. The value without IbpA was set to 1. Statistical analysis: Error bars represent SD*; n* = 3 biological replicates. The statistical significance of differences was assessed using the Student’s *t*-test (\**p* < 0.05, \*\**p* < 0.01).

### The control of the σ^32^ levels by IbpA is independent of the degradation pathway through DnaK

Chaperone-mediated degradation of σ^32^ is one mechanism for regulating its abundance (3, 12) (Fig. 4A). To examine whether IbpA affects σ^32^ degradation, we investigated whether IbpA’s effect on σ^32^ is abolished in cells with deleted DnaK or the σ^32^ I54N mutation, which are known to inhibit the degradation pathway (2, 3, 6, 12–14, 20, 35). We evaluated the endogenous σ^32^ expression in an *E. coli* strain that deleted the *dnaK*-*dnaJ* operon (Δ*dnaKJ*), with or without the IbpA overexpression. Our data showed that even in the Δ*dnaKJ* cells, IbpA overexpression significantly suppressed σ^32^ expression (Fig. 4B), suggesting that IbpA-mediated σ^32^ suppression occurs independently of the DnaK-mediated degradation pathway. In addition, we found that DnaK/DnaJ supplementation in the IbpA-overexpressing Δ*dnaKJ* strain further decreased the σ^32^ level, indicating that the effects of IbpA and DnaK are additive. We next used the σ^32^ I54N mutant, which is defective in chaperone-mediated degradation and inactivation (12, 15–17). The σ^32^ I54N mutant expression was upregulated by the *ibpA-ibpB* deletion and decreased by IbpA overexpression (Fig. 4C). These results demonstrate that IbpA affects the σ^32^ level at the translational level but not at the degradation step.

**Fig 4.**
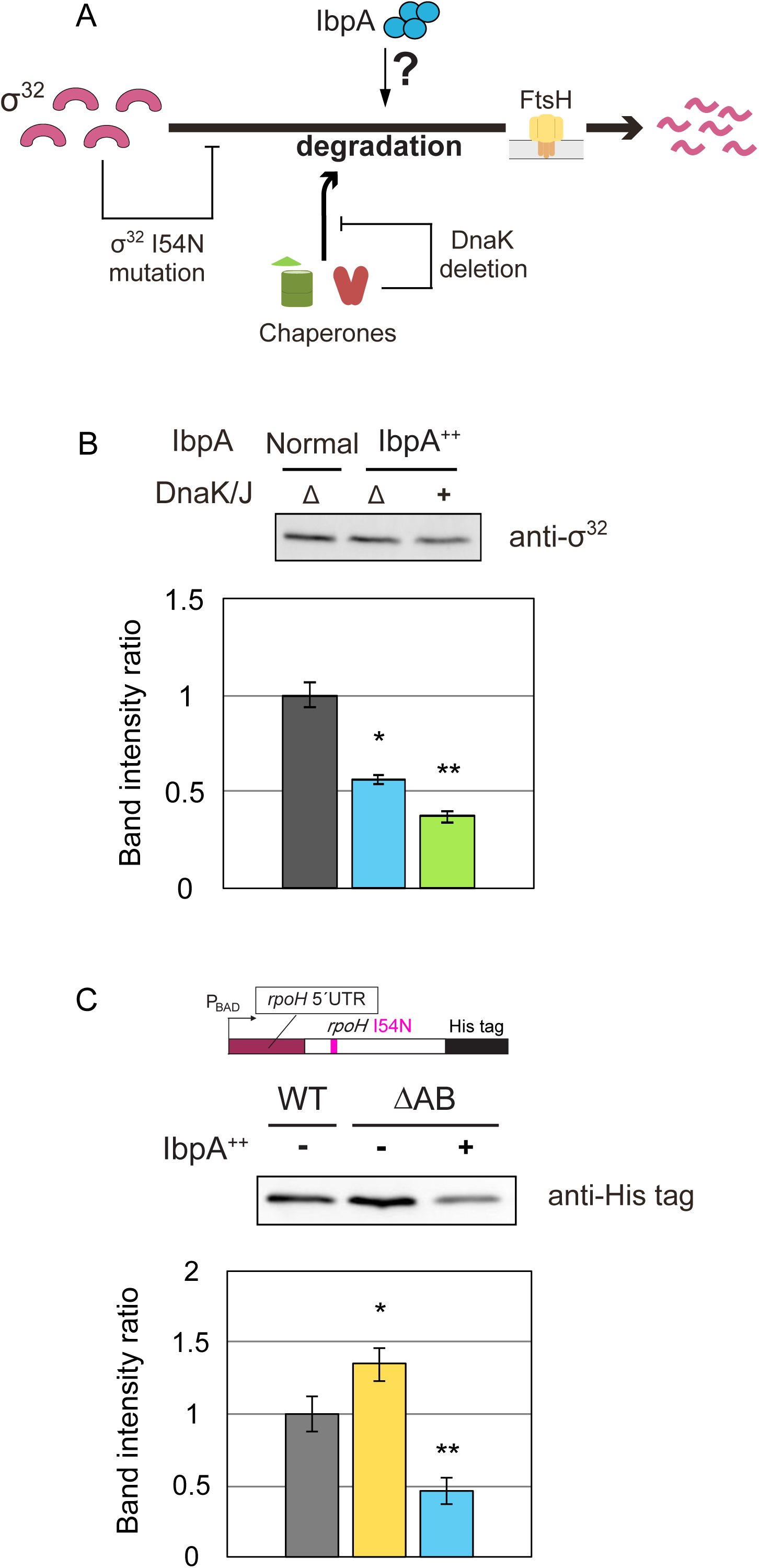
IbpA-mediated control of the σ^32^ level is independent of degradation via the DnaK pathway. (A) Model illustrating regulation of σ^32^ degradation. (*B*) Western blotting used to evaluate endogenous σ^32^ levels in the *dnaK-dnaJ* operon deleted strain (Δ*dnaKJ*) with or without IbpA and DnaK/DnaJ expression. Normal, *E. coli* wild-type (*i.e.* endogenous level of IbpA). IbpA^++^, *E. coli* wild-type overexpressing IbpA from a plasmid. Δ, Δ*dnaKJ* strain. +, Δ*dnaKJ* strain complemented with DnaK/DnaJ by expression from a plasmid. Anti-σ^32^ antibody was used for detection. Relative band intensities of three biological replicates are shown below. The value in the Δ*dnaKJ* strain was set to 1. (*C*) Western blotting evaluating the effect of the σ^32^ I54N mutation using the reporter system shown in Fig. 3B. Anti-His tag antibody was used for detection. Relative band intensities of three biological replicates are shown below. The value in the *E. coli* wild-type was set to 1. Statistical analysis: Error bars represent SD. The statistical significance of differences was assessed using the Student’s *t*-test (\**p* < 0.05, \*\**p* < 0.01).

### IbpA contributes to the shut-off of σ^32^ during the heat shock response

It is widely acknowledged that σ^32^ levels experience a temporary increase during heat shock followed by a decrease, a phase referred to as “shut-off”, which is critical for the recovery to normal conditions (3, 6, 7). We tested whether the IbpA-mediated regulation of σ^32^ affects the shut-off phase. In order to determine the level shift of σ^32^ accompanied by heat shock, we assessed the abundance of endogenous σ^32^, which was normalized by that of FtsZ, a protein expressed constitutively. In wild-type *E. coli* cells, the σ^32^ level was at its maximum 5 min after the onset of heat stress and subsequently decreased to a normal level 10 min later (Fig. 5A), consistent with previous studies (3, 7). Upon exposure to heat stress in the Δ*ibpAB* strain, the σ^32^ level increased similarly to the wild-type strain. However, in contrast to the wild-type, the increased level of σ^32^ continued even at 10 min and was only shut-off to the normal level after 30 min (Fig. 5A). Unlike the wild-type and the Δ*ibpAB* strains, overexpression of IbpA significantly suppressed σ^32^ expression and heat stress-response of σ^32^ (Fig. 5A). Along with the change in the amount of σ^32^ in *E. coli* with varying IbpA levels, the amount of Hsps also affected. For example, the amounts of GroES increased or decreased in the *E. coli* with the depletion or the overexpression of IbpA, respectively (Fig. S4). These findings demonstrate that the overexpression or loss of IbpA perturbs the normal heat shock response.

**Fig 5.**
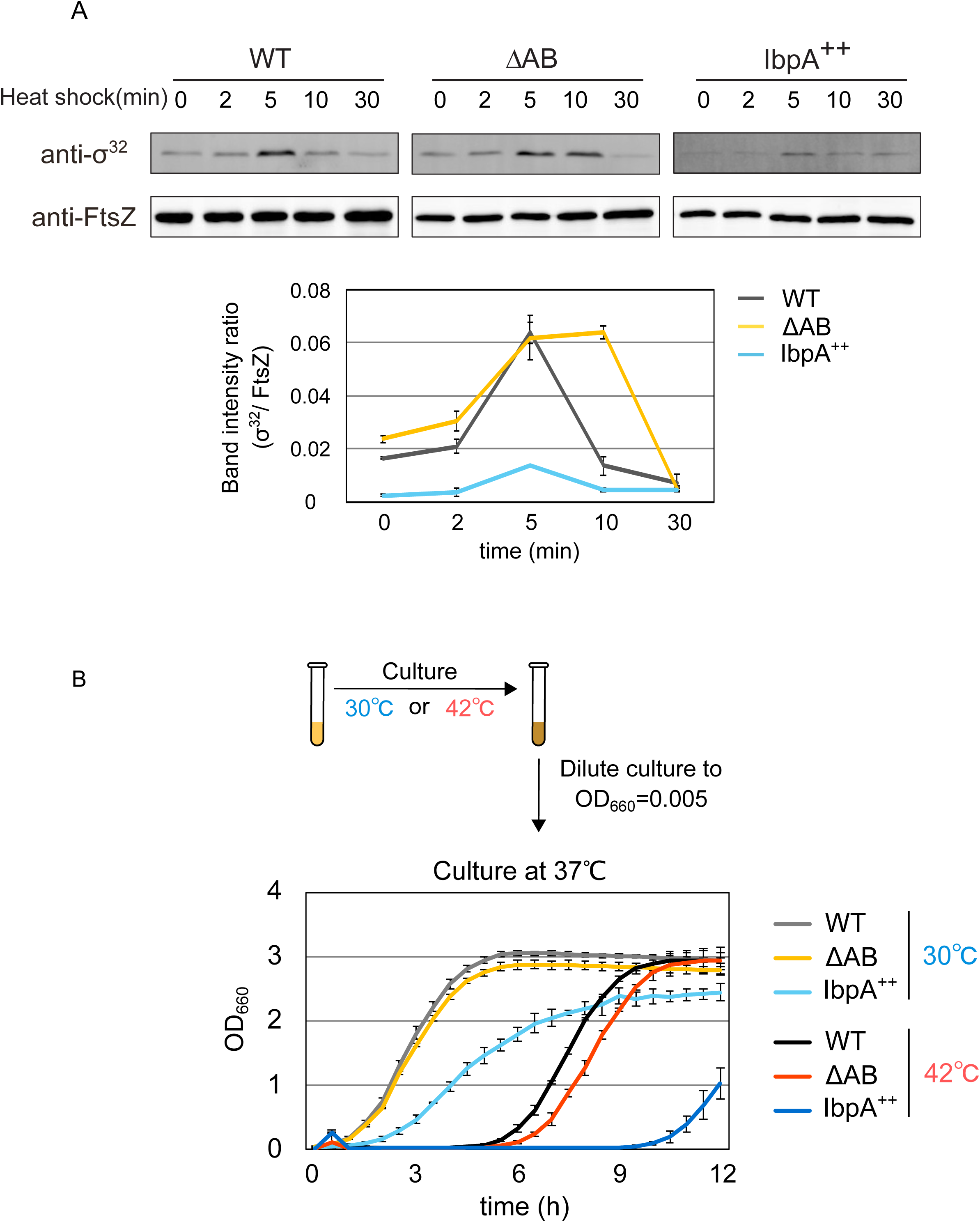
IbpA contributes to the shut-off of σ^32^ during heat shock response. (*A*) Evaluation of the σ^32^ during heat shock response. *Top*: Western blotting of endogenous σ^32^ and FtsZ in wild-type (WT), *ibpAB* deleted (Δ*AB*), and Δ*AB* cells overexpressing IbpA (IbpA^++^,) cells. Anti-σ^32^ and anti-FtsZ antibodies were used to detect σ^32^ and FtsZ, respectively. FtsZ, a constitutively expressed protein, was used to normalize the band intensity of σ^32^. HS (min): the minutes from the start of heat shock. *Bottom*: Quantification of σ^32^ levels normalized by FtsZ levels. Error bars represent SD; *n* = 3 biological replicates. (*B*) The growth curve of *E. coli* after recovery from heat shock. The cells were grown at 30 or 42 °C for 2 h and then diluted to allow growth at 37 °C. WT: wild-type cells; *ΔAB*: *ibpAB* deleted strain; *IbpA*^++^: Δ*AB* cells overexpressing IbpA. Error bars represent SD; *n* = 3 biological replicates.

Given that the shut-off phase is associated with the recovery from heat stress, we next examined the effect of IbpA on cell growth after exposure to heat shock at 42 °C for 2 hours. The cell growth rate during recovery after stress was slightly, but statistically significantly, impaired in the Δ*ibpAB* strain compared to the wild strain (Fig. 5B). In the IbpA-overexpression strain, which exhibited slower growth than the wild strain even at 37 °C as previously shown (27), the recovery after stress was extremely poor (Fig. 5B).

## Discussion

IbpA, sHsp in *E. coli*, has been identified as a chaperone that sequesters aggregation proteins by co-aggregation during stress. Recently, it has also been shown to act as a mediator for self-regulation at the posttranscriptional level (27). This study provides compelling evidence that IbpA represses the translation of *rpoH* mRNA, which is a novel target for IbpA-mediated regulation in *trans*, since previously known targets were limited to *ibpA* and *ibpB* mRNAs. The ability of IbpA to regulate other ORF mRNAs suggests that its effect on σ^32^ could influence the heat shock response, in addition to the FtsH-mediated degradation assisted by chaperones like DnaK and GroEL, which are already well-established.

We have found that the abundance of oligomeric IbpA influences the level of σ^32^ in a manner that is dependent on the *rpoH* 5’ UTR. Additionally, we reconstituted the suppression of translation by IbpA in the PURE system. These findings strongly suggest that the fundamental mechanism of translation suppression by IbpA on *rpoH* is comparable to that on *ibpA* or *ibpB*. Since the translation of *ibpA* and *rpoH* mRNAs is governed by thermoresponsive mRNA secondary structures (RNATs), IbpA oligomers could attach to the 5’ UTR of *rpoH* mRNA to repress translation under normal conditions. We previously proposed a titration model for the self-regulation of IbpA during heat stress. In this model, IbpA-mediated self-repression of translation via mRNA binding is abolished by the recruitment of IbpA to protein aggregates (27). The role of IbpA as a regulator of its own expression via sensing of protein aggregation may be relevant to the control of σ^32^. This mechanism is reasonable given the function of sHsps as the first line of defense against aggregation stress.

Then, we propose a model for the regulatory role of IbpA in the σ^32^ response (Fig. 6). In non-stress conditions, IbpA suppresses *rpoH* translation, while under heat stress conditions, IbpA acts as a “sequestrase” chaperone to recruit aggregation-prone proteins. This titrates the free IbpA away to alleviate the translation suppression. Concurrently, the high temperature-induced disruption of RNAT in the *rpoH* 5’ UTR enhances σ^32^ expression. The titration mechanism occurs at the endogenous level of IbpA, as artificial overexpression of IbpA fully inhibits σ^32^ translation, thereby compromising the heat shock response (*cf.* Fig. 2E, Fig. 5A).

**Fig 6.**
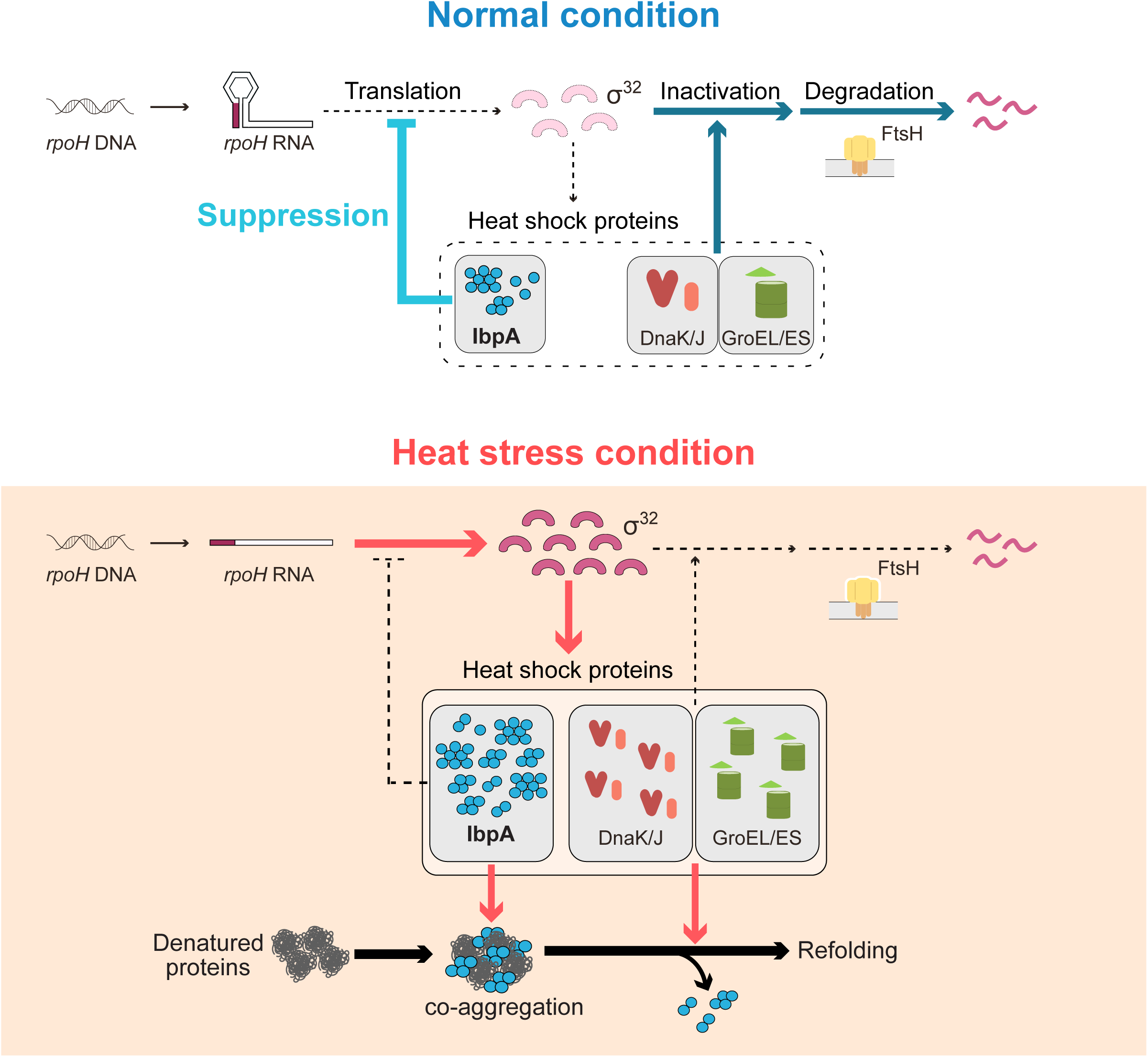
Model of the IbpA’s contribution in regulating the heat shock response. *Top*: Under normal conditions, σ^32^ is inactivated by DnaK/J and GroEL/ES, and then degraded by FtsH at posttranslational level. In addition, *rpoH* translation is suppressed by a thermoresponsive mRNA secondary structure (RNAT) and IbpA. *Bottom*: Under stress conditions, chaperones are recruited into heat-denatured proteins, leading to the release of σ^32^ repression due to inactivation/degradation and translational repression. This results in an increased functional σ^32^ level, eventually upregulating Hsps.

A significant question regarding the novel function of IbpA concerns the mechanism by which IbpA interacts with various types of mRNAs to inhibit translation, given the lack of putative RNA-binding motif in IbpA. Our previous findings indicated that oligomeric IbpA state and secondary structures in RNAT are critical to translation suppression (27). Thus, the complex interplay between IbpA and mRNAs might be essential to its suppression ability. Moreover, understanding the difference between IbpA and IbpB, which lacks suppression activity, could provide valuable insight into this issue.

IbpA-mediated regulation of σ^32^ using a titration mechanism is conceptually analogous to the well-known σ^32^ regulatory mechanism involving DnaK/J, GroEL/ES, and FtsH (2, 3, 6, 12–14). Interestingly, the IbpA-dependent regulation is independent of the DnaK-mediated pathway, as the IbpA level affects the amount of σ^32^ even in the DnaK-deleted strain. How does this IbpA-mediated regulation relate to the regulation involving DnaK and other factors? The primary difference lies in the level of control, whether at the post-translational or translational level. The DnaK/J and GroEL/ES chaperone systems suppress σ^32^ activity under normal conditions and facilitate FtsH-dependent degradation of σ^32^ (3, 11, 12). This inactivation-degradation mechanism relies on pre-existing σ^32^ proteins after translation. In contrast, the IbpA-mediated mechanism operates at the translational level and can rapidly induce σ^32^ expression upon heat stress. Therefore, combining the two-layered mechanism at both translational and post-translational levels would be complementary, resulting in a more robust and flexible induction of σ^32^ during stress. Additionally, previous reports have indicated that excess Hsp and σ^32^ adversely affect growth (11, 15), further highlighting the importance of the two-layered mechanism.

In the pathway mediated by DnaK-GroEL, regulation of σ^32^ occurs not only in degradation but also in activity, with DnaK/J and GroEL binding to the inactive state of σ^32^ (3, 12). While our study focuses solely on IbpA’s regulation of *rpoH* translation, it is plausible that IbpA, acting as a chaperone, may also contribute to regulating σ^32^ activity.

In a previous study, we proposed that the IbpA-mediated self-repression is a “safety catch” mechanism that tightly regulates IbpA expression, which can be harmful under normal conditions. The RNAT structure of *ibpA* is vital for its regulation, and its RNAT can partially open even under normal conditions, such as 37 °C (25, 27). This study also indicates that the *rpoH* 5’ UTR, responsible for the structure of RNAT, is required for translational regulation. The RNAT of *rpoH* opens between 30-40 °C, permitting partial translation of *rpoH* (22). IbpA may act as a “safety catch” for the *rpoH* RNAT to prevent the unnecessary expression of σ^32^ under conditions without stress. Indeed, since excess Hsp and σ^32^ have a detrimental effect on growth (11, 15), IbpA-mediated tight repression of σ^32^ under normal conditions is reasonable. The previously-known role of IbpA is to sequester aggregation upon stress (35–38), in contrast to the essential housekeeping roles of DnaK/J and GroEL/ES, even under normal conditions. Therefore, IbpA, without an apparent housekeeping role, may be suitable to function as a safety catch to prevent the production of unnecessary amounts of Hsps under normal conditions.

Our results regarding IbpA have implications for the σ^32^ shut-off phase during heat stress. Although the shut-off mechanism depends on the accelerated degradation mediated by the excessive amount of chaperones produced during heat shock (3, 6, 7), the level of σ^32^ is shut-off even when the degradation pathway is inhibited (18, 19) or with the hyperactive σ^32^ I54N mutant (17), suggesting an alternative shut-off mechanism. Since IbpA regulates the level of σ^32^ at a translational level, a different level from the canonical chaperone-mediated degradation regulation, IbpA may be responsible for regulating σ^32^ synthesis during the shut-off phase. Indeed, in the Δ*ibpAB* strain, the σ^32^ shut-off, which occurs 10 min after the initiation of heat shock in the wild-type strain, does not occur but shuts off after 30 min (Fig. 5A). The σ^32^ shut-off observed at 30 min in the Δ*ibpAB* cells would be induced by the canonical degradation pathway. The delayed shut-off in the Δ*ibpAB* cells would be interpreted that IbpA affects the initial phase of the shut-off, followed by FtsH-mediated degradation assisted by induced chaperones such as DnaK (Fig. 6).

We observed that the Δ*ibpAB* strain exhibited a slight delay in the recovery of growth after exposure to a heat shock at 42 °C. This observation supports a previous finding that the Δ*ibpAB* cells display a delayed resolubilization of protein aggregates after heat shock (39), which may result from a slightly delayed recovery from stress in the Δ*ibpAB* cells (Fig. 5B). Furthermore, it is known that the deletion of the *ibpA-ibpB* operon increases susceptibility to chaperone depletion (39). However, while IbpA can compensate for this depletion, its paralog IbpB cannot (33). We speculate that this difference arises from the subtle but significant difference between IbpA and IbpB. Besides their differences in chaperone function (33), the exclusive role of IbpA in suppressing the translation of σ^32^ might correlate with the phenotypes observed in the Δ*ibpAB* strain.

What is the physiological significance of IbpA-mediated suppression of *rpoH* translation? One possibility is that the regulation of expression at the translation level is generally rapid and thus well-suited for immediate induction upon heat shock. Another advantage of tight suppression of the σ^32^ levels by IbpA under normal conditions is the suppression of Hsp expression, which is unnecessary in the absence of stress. In addition, the cost of ribosome occupancy for the translation of Hsps (5) would be reduced.

Lastly, are there other targets for IbpA-mediated translation suppression? Our findings suggest that IbpA overexpression decreases the level of endogenous *rpoH* mRNA without affecting its degradation rate (Fig. S3A, B). This result suggests that certain factors affect *rpoH* transcription in cells with IbpA-overexpression. Indeed, proteomics analysis has shown that expression of genes controlled by CRP, the dual transcriptional regulator of *rpoH*, is decreased in cells with IbpA overexpression (Supplementary Dataset S2) (40). Further analysis is necessary to elucidate the complete regulatory picture mediated by IbpA.

## Materials and Methods

### E. coli strains

Table S1 provides a comprehensive list of *E. coli* strains employed in this study. DH5α strain was used for cloning. The BW25113 strain was used for each assay. The chromosomal *ibpA-ibpB* or *dnaK-dnaJ* operons were deleted via previously described procedures (41). The DNA fragment amplified from JW3664 (Δ*ibpA*::FRT-Km-FRT) (42), using the primers PT0456 and PM0195, and that from JW3663 (Δ*ibpB*::FRT-Km-FRT) (42), using the primers PT0457 and PM0196, were annealed and amplified together using PT0456 and PT0457. The purified DNA was electroporated into the *E. coli* strain BW25113 harboring pKD46, and the transformant that was kanamycin-resistant at 40 µg/mL was stored as BW25113Δ*ibpAB*. A list of primers utilized in the study is available in Table S2.

### Plasmids

Plasmids were constructed using standard cloning procedures and Gibson assembly. Plasmids designed for overexpression purposes, including pCA24N-*ibpA*, pCA24N-*ibpA*-*AEA*, and pCA24N-*gfp*, were constructed using DNA fragments that were amplified from pCA24N (43), superfolder GFP (44, 45), or *E. coli* genomic DNA. Plasmids were also constructed for the evaluation of expression, including pBAD30-*dnaK*promoter-*gfp*, pBAD30-*groES*promoter-*gfp*, pBAD30-*dnaK* 5’ UTR-*gfp*, pBAD30-*groES* 5’ UTR-*gfp*, pBAD30-*rpoH* 5’ UTR-*rpoH*, pBAD30-*rpoH*, pBAD30-*rpoH* 5’ UTR-*rpoH-I54N*, and pBAD30-*ibpAB*operon. These plasmids were constructed using DNA fragments amplified from pBAD30 (46), superfolder GFP (45), derived from a previously constructed plasmid (44), and *E. coli* genomic DNA. pKJE7 (Takara) was used to supply DnaKJ to the *dnaK-dnaJ* deleted cells. The primers employed in the cloning process are listed in Table S2.

### MS method

*E. coli* BW25113 cells harboring a plasmid carrying the IbpA or GFP genes were grown at 37 °C in LB medium. The plasmids were induced with 50 mM isopropyl-β-D-thiogalactopyranoside (IPTG) at an OD_660_ of 0.5. Cells were collected during the exponential phase (∼1.0 Abs_660_) by centrifugation at 20,000 ×*g*. Sample preparation for liquid chromatography (LC)-MS/MS was conducted in accordance with a previous study with some modifications (47, 48). The quantitative LC-MS/MS analysis was conducted by using SWATH (sequential windowed acquisition of all theoretical mass spectra)-MS acquisition (49). All LC-MS/MS measurements were performed with Eksigent nanoLC 415 and TripleTOF 6600 mass spectrometer (AB Sciex, U.S.A.). The trap column used for nanoLC was a 5.0 mm × 0.3 mm ODS column (L-column2, CERI, Japan), and the separation column was a 12.5 cm × 75 μm capillary column packed with 3 μm C18-silica particles (Nikkyo Technos, Japan). The libraries for SWATH acquisition were constructed on the basis of in-house IDA (information-dependent acquisition) measurements.

The SWATH acquisition and data analysis procedures were performed according to the previous study (47). Only proteins detected in all three measurements for both samples were used for calculating fold changes. The resulting protein intensities were averaged using an in-house R script. The *p*-value was determined using Welch’s *t*-test and corrected using the Benjamini-Hochberg method for multiple comparisons, employing the “p.adjust” function in R (for Windows, version 4.1.2). Gene ontology (GO) enrichment analysis for increased/decreased proteins was performed with DAVID (50), using all detected protein data as background for enrichment analysis.

### SDS-PAGE and western blotting

*E. coli* BW25113 cells harboring a plasmid for evaluation were precultured at 30 °C for 16 h in LB medium. The cells were then grown to an OD_660_ of 0.4∼0.6 in LB medium at 30 °C. For the pBAD30 plasmid, 2 × 10^-4^ % arabinose was used for induction of the reporter, while 0.1 mM IPTG was added 3 h after the start of the culture for pCA24N. In the co-expression assay, *E. coli* BW25113 cells harboring plasmids for evaluation and pCA24N plasmids were used. The induction of protein co-expression was performed with 0.1 mM IPTG, added 2 h after starting the culture. Cell cultures were harvested and mixed with an equal volume of 10% trichloroacetic acid, to terminate the biological reactions and precipitate macromolecules. After standing on ice for at least 15 min, the samples were centrifuged at 20,000 ×*g* for 3 min at 4 °C, and the supernatant was removed by aspiration. Precipitates were washed with 1 mL of acetone by vigorous mixing, centrifuged again, and dissolved in 1× SDS sample buffer (62.5 mM Tris-HCl, pH 6.8, 5% 2-mercaptoethanol, 2% SDS, 5% sucrose, 0.005% bromophenol blue) by vortexing for 15 min at 37 °C. The samples were separated by SDS-PAGE, transferred to PVDF membranes, and blocked using 5% non-fat milk in Tris-buffered saline with 0.002% Tween-20. Mouse anti-sera against GFP (mFx75, Wako), mouse anti-sera against His-tag (9C11, Wako), rabbit anti-sera against DnaK (abcam), rabbit anti-sera against GroES (a gift from Dr. Ayumi Koike-Takeshita at Kanagawa Institute of Technology), rabbit anti-sera against σ^32^ (Biolegend), and rabbit anti-sera against FtsZ (a gift from Dr. Shinya Sugimoto at Jikei Medical University) were used as primary antibodies at a 1:10,000 dilution. Secondary antibodies, Cy3 conjugated anti-mouse IgG (Invitrogen) and Cy5 conjugated anti-rabbit IgG (Abcam) were also used. Blotted membranes were detected with a fluorescence imager (Amersham Typhoon RGB, Cytiva), and band intensities were quantified with analysis software (Image quant, Cytiva).

### Quantitative RT-PCR

To quantify mRNA relative amounts, *E. coli* BW25113 cells harboring a plasmid harboring the reporter gene were precultured at 30 °C for 16 h in LB medium. The cells were then grown in 5 ml of LB medium at 37 °C to an OD_660_ of 0.4∼0.6. The reporter gene carried in pBAD30 plasmid was induced with 2 × 10^-4^ % arabinose, and protein co-expression was induced by adding 0.1 mM IPTG 2 h after initiating the culture. The cultures were pelleted at 10,000 ×*g* for 3 min at 4 °C.

For quantifying mRNA degradation, *E. coli* BW25113 cells harboring a plasmid carrying the reporter gene were precultured at 30 °C for 16 h in LB medium. The cells were then grown in 20 ml of LB medium at 30 °C to an OD_660_ of 0.4∼0.6. The reporter gene in the pBAD30 plasmid was induced with 2 × 10^-4^ % arabinose. After 2.5 h from of culture initiation, cells were aliquoted into 5 mL portions and treated with 250 µM Rifampicin. Cultures were sampled at 2, 5, and 10 min after Rifampicin treatment, and the cells were pelleted at 10,000 ×*g* for 3 min at 4 °C.

Total mRNA was extracted using Tripure Isolation Reagent (Merck) and treated with recombinant DNase I (Takara). The treated RNA was subsequently purified using an RNeasy Mini kit (Qiagen). The samples were prepared using a Luna Universal One-Step RT-qPCR Kit (New England Biolabs). Quantitative RT-PCR was performed with an Mx3000P qPCR system (Agilent) and analyzed using the MxPro QPCR software (Agilent). The ΔΔCt method (51) was used to normalize the amount of target mRNA with the *ftsZ* mRNA. The primers used for PCR are listed in Table S2.

### Cell-free translation

The transcription–translation-coupled PURE system (PUREfrex, GeneFrontier) reaction was carried out at 37 °C for 2 h in the presence or absence of 1 µM IbpA, which included Cy5 labeled tRNA^fMet^. After protein synthesis, SDS-sample buffer (0.125 M Tris-HCl (pH 6.8), 10% (v/v) 2-mercaptoethanol, 4% (w/v) SDS, 10% (w/v) sucrose, 0.01% (w/v) bromophenol blue) was added and incubated at 95 °C for 5 min. The samples were then separated by SDS-PAGE and detected using a fluorescence imager (Amersham Typhoon RGB, Cytiva) at the 633nm wavelength. The band intensity was quantified with image analysis software (Image quant, Cytiva).

### Statistical Analysis

Student’s *t*-test was used for calculating statistical significance, with a two-tailed distribution with unequal variance. All experiments were conducted at least three times independently, and the mean values ± standard deviation (SD) were represented in the figures.

## Acknowledgments

We thank Tatsuya Niwa for technical support on the LC-MS/MS analysis; the Bio-support Center at Tokyo Tech for DNA sequencing; the Cell Biology Center Research Core Facility at Tokyo Tech for the Q-Exactive and TripleTOF 6600 mass spectrometry measurements; Ayumi Koike-Takeshita for the anti-GroES antibody; and Shinya Sugimoto for the anti-FtsZ antibody. This work was supported by MEXT Grants-in-Aid for Scientific Research (Grant Numbers JP26116002, JP18H03984, and JP20H05925 to HT, JP21K20631, and JP22K14860 to TM).

## Author Contributions

T.M. performed experiments; T.M. and H.T. conceived the study, designed experiments, and analyzed the results; H.T. supervised the entire project; T.M. and H.T. wrote the manuscript.

## Classification

Biological sciences> Biochemistry

