## supplemental figures and tables for "The *Escherichia coli* small heat shock protein IbpA plays a role in regulating the heat shock response by controlling the translation of σ^32^"

**This PDF file includes:**

Supplementary Figures S1 to S4

Supplementary Tables S1-S3

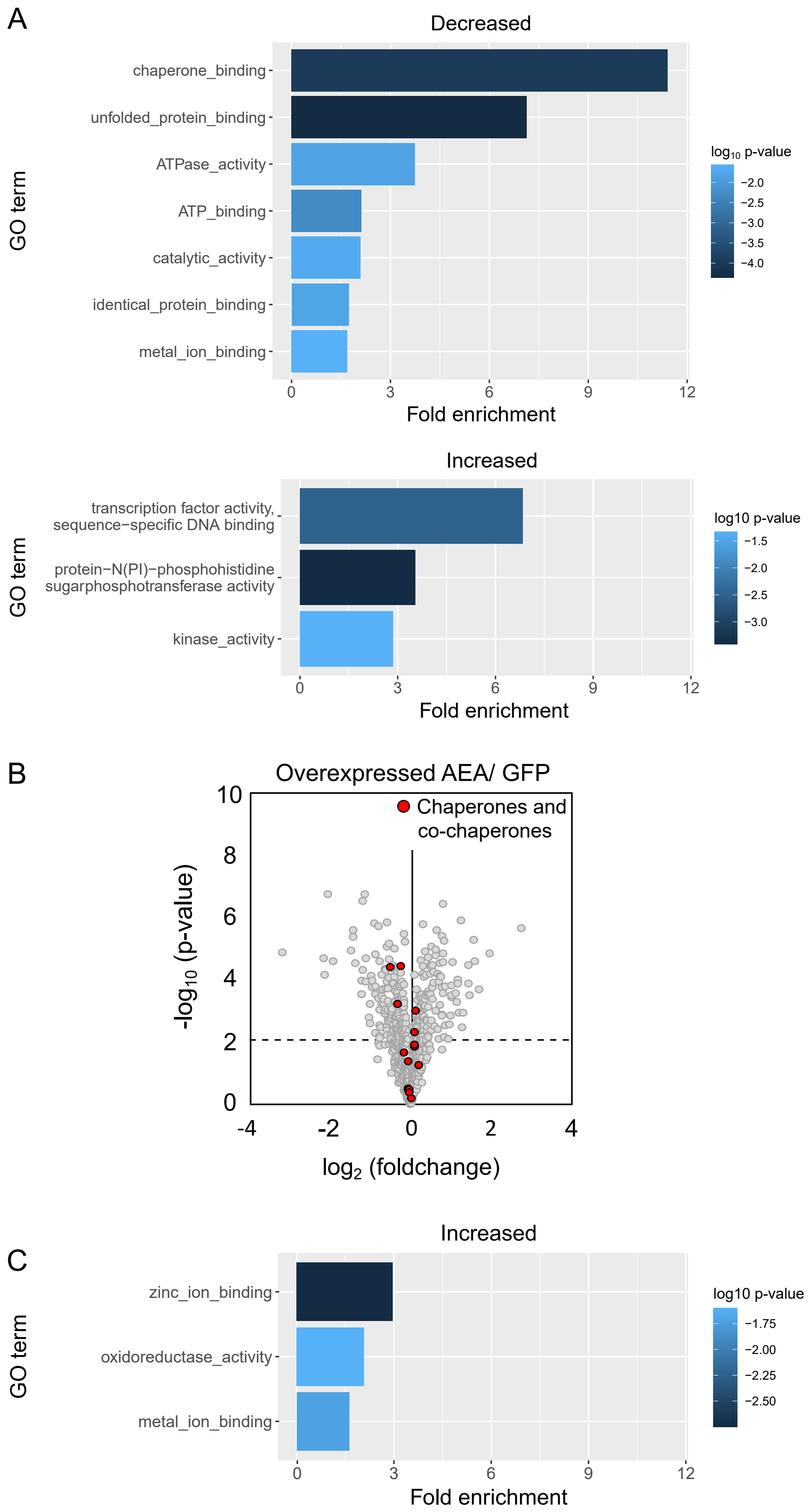
A
Decreased
chaperone_binding
unfolded_protein_binding
ATPase_activity
ATP_binding
catalytic_activity
identical_protein_binding
metal_ion_binding
GO term
Fold enrichment
log10 p-value
Increased
transcription factor activity, sequence−specific DNA binding
protein−N(PI)−phosphohistidine sugarphosphotransferase activity
kinase_activity
B
Overexpressed AEA/ GFP
Chaperones and co-chaperones
-log10 (p-value)
log2 (foldchange)
C
zinc_ion_binding
oxidoreductase_activity
metal_ion_binding

**Fig. S1. Enrichment of heat shock proteins in the group exhibiting decreased expression upon overexpression of IbpA.**

(*A*) Comparative proteomics analysis was conducted on the *E. coli* wild-type strain overexpressing IbpA and GFP to investigate the proteins that were either reduced or increased by IbpA overexpression through Gene Ontology (GO) analysis. The fold enrichment of each GO term is represented on the horizontal axis, while the *p*-value is by the color of each bar, with the darker colors indicating higher statistical significance. (*B*) The volcano plot displays the protein expression ratio between the IbpA-AEA mutant and GFP overexpression in *E. coli* cells. Each dot in the plot represents the fold change and *p*-value of each protein identified through the comparative proteomics analysis. The dashed line indicates a *p*-value of 0.01. Proteins categorized as “chaperone binding” and “unfolded protein binding” in the GO analysis are represented by red dots. (*C*) GO analysis was conducted on the proteins increased by IbpA-AEA overexpression in *E. coli* cells using comparative proteomics analysis. Enriched GO terms for the proteins increased by the IbpA-AEA overexpression are shown, with the same display format as in (*A*). Note that there were no enriched GO terms with a high statistical significance for the proteins decreased by IbpA-AEA overexpression.

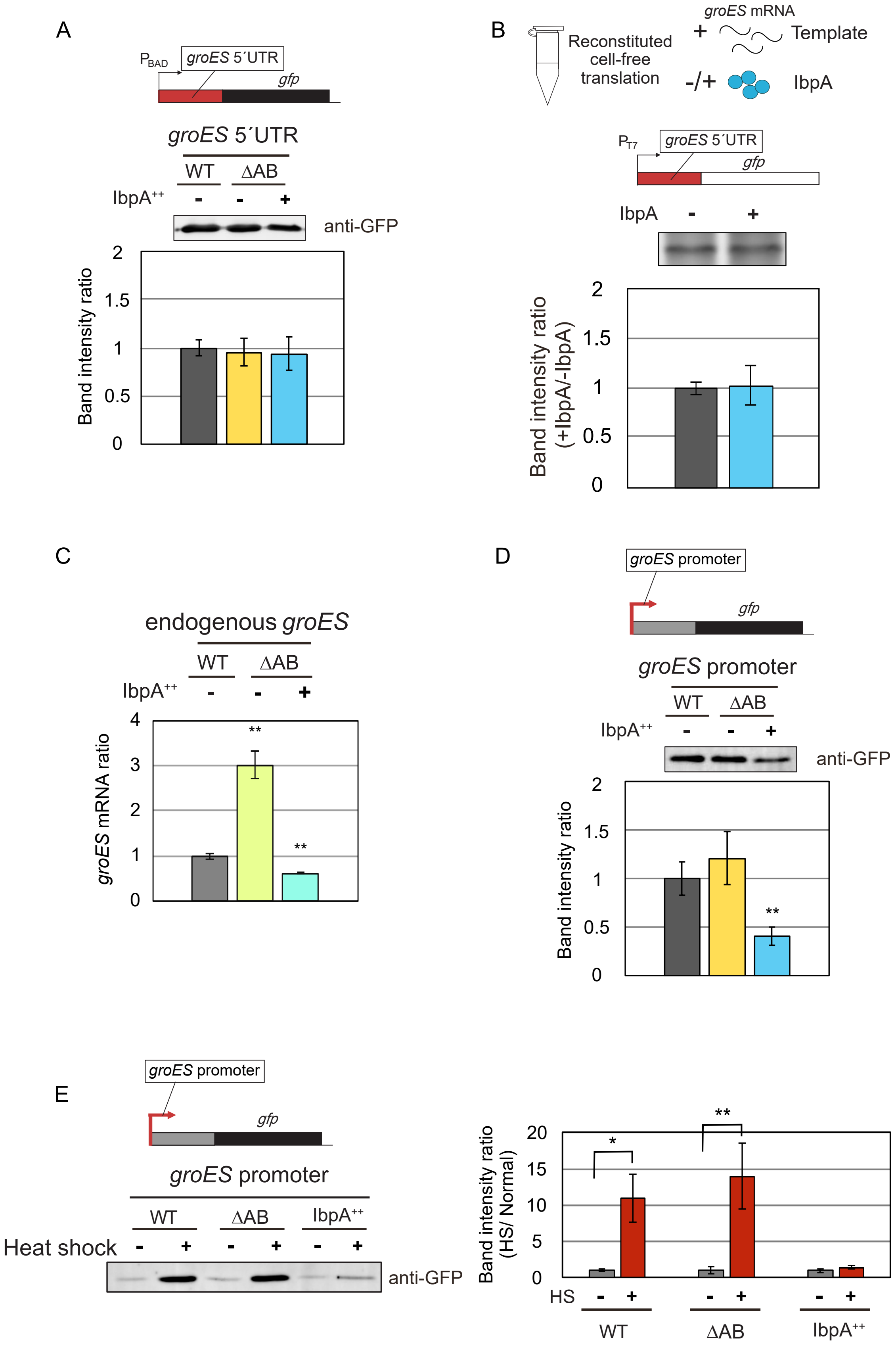
A
PBAD
groES 5´UTR
gfp
groES 5´UTR
WT
ΔAB
IbpA++
-
-
+
anti-GFP
Band intensity ratio
2
1.5
1
0.5
0
B
groES mRNA
Reconstituted cell-free translation
+
Template
-/+
IbpA
PT7
groES 5´UTR
gfp
IbpA
-
+
Band intensity ratio (+IbpA/-IbpA)
2
1.5
1
0.5
0
C
endogenous groES
WT
ΔAB
IbpA++
-
-
+
groES mRNA ratio
4
3
2
1
0
**
**
D
groES promoter
gfp
groES promoter
WT
ΔAB
IbpA++
-
-
+
anti-GFP
Band intensity ratio
2
1.5
1
0.5
0
**
E
groES promoter
gfp
groES promoter
WT
ΔAB
IbpA++
Heat shock
-
+
-
+
-
+
anti-GFP
Band intensity ratio (HS/ Normal)
20
15
10
5
0
*
**
HS
-
+
-
+
-
+
WT
ΔAB
IbpA++

**Fig. S2. IbpA suppresses the expression of GroES at a transcription level.**
(*A*) Western blotting of GFP harboring *groES* 5′ UTR in *E. coli* wild-type strain (WT) or the *ibpAB* operon-deleted strain (Δ*AB*). $IbpA^{++}$, *E. coli* Δ*AB* cells overexpressing IbpA. The relative band intensities of three biological replicates are shown below. The value in the *E. coli* wild-type was normalized to 1. (*B*) Cell-free translation of the *gfp* reporter was carried out in the absence (-) or the presence (+) of purified IbpA using a reconstituted protein synthesis system (PURE system). The value without IbpA was set as 1. (*C*) Evaluation of the endogenous mRNA level of *groES* by qRT-PCR in wild-type cells (*left*), Δ*AB* cells (*middle*), and Δ*AB* cells expressing IbpA (*right*). The value in the *E. coli* wild-type was set to 1. (*D*) Western blotting of GFP expressed by the endogenous *groES* promoter in *E. coli* cells. The value in the *E. coli* wild-type was set to 1. (*E*) Western blotting (*left*) and the relative band intensities (*right*) of GFP expressed by the endogenous *groES* promoter in *E. coli* cells with (*red*) or without (*gray*) heat shock (HS). The values without heat shock were set to 1.
Statistical analysis for (*A*)-(*E*): Error bars indicate the standard deviation (SD); $n$ = 3 biological replicates. Student's *t*-test was employed to assess the statistical significance of differences ($*p < 0.05$, $**p < 0.01$).

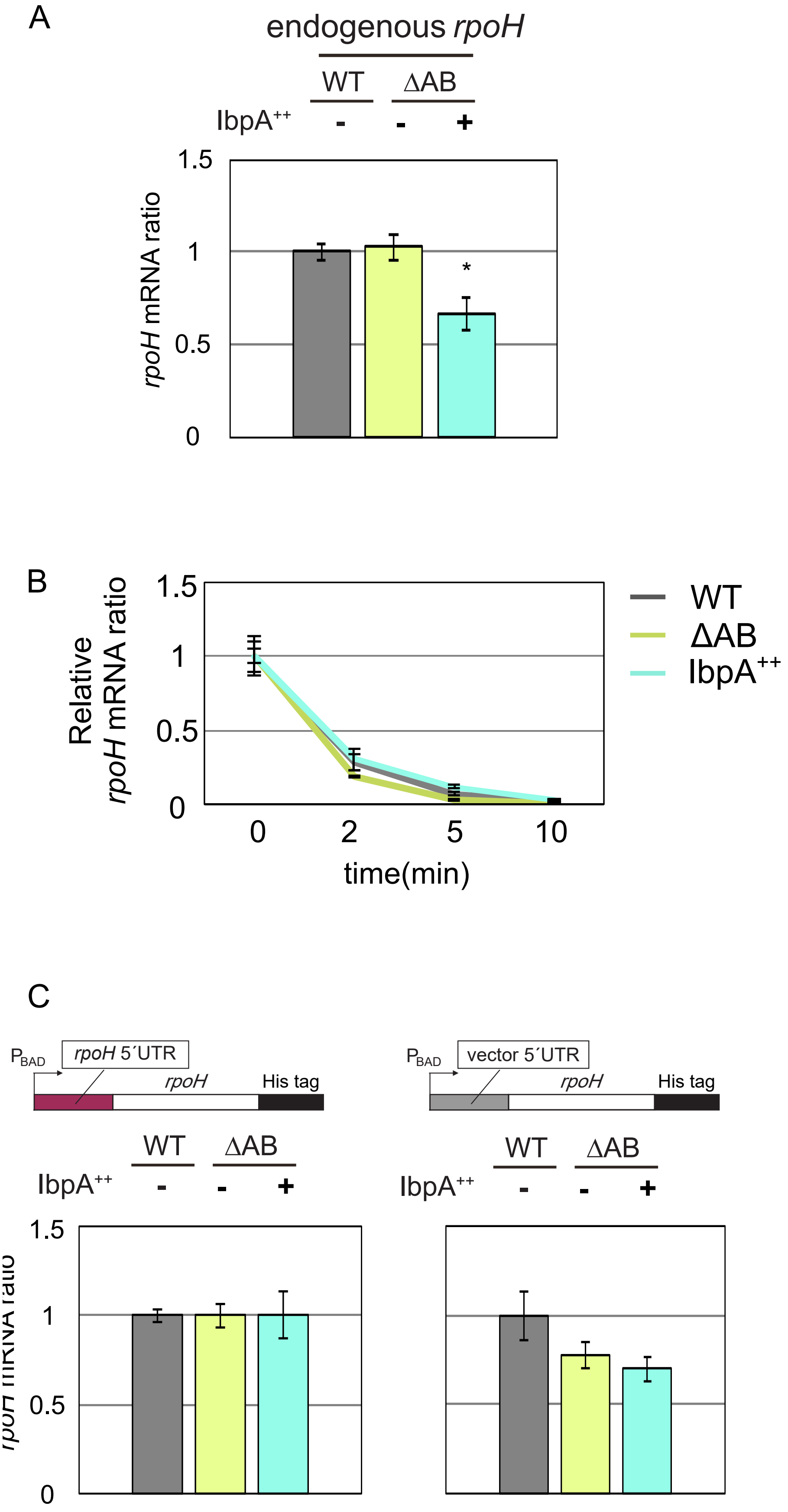
A
endogenous rpoH
WT
ΔAB
IbpA++
-
-
+
rpoH mRNA ratio
1.5
1
0.5
0
*
B
Relative rpoH mRNA ratio
1.5
1
0.5
0
0
2
5
10
time(min)
WT
ΔAB
IbpA++
C
PBAD
rpoH 5´UTR
rpoH
His tag
PBAD
vector 5´UTR
rpoH
His tag
WT
ΔAB
IbpA++
-
-
+
WT
ΔAB
IbpA++
-
-
+
1.5
1
0.5
0

**Fig. S3. IbpA suppresses the $\sigma^{32}$ expression independent of transcriptional or degradation controls.**

(*A*) Quantification of endogenous *rpoH* mRNA levels by qRT-PCR in *E. coli* wild-type strain (WT), *ibpAB* operon-deleted strain (Δ*AB*), and *E. coli* Δ*AB* cells overexpressing IbpA (IbpA$^{++}$). The value in *E. coli* wild-type was normalized to 1. (*B*) Degradation kinetics of endogenous *rpoH* mRNA in *E. coli*. Time is represented in minutes from the addition of Rifampicin to inhibit RNA polymerases. The value at time 0 was set to 1. (*C*) Quantification of the levels of *rpoH* mRNA harboring the 5' UTR of *rpoH* or a vector under the control of an arabinose promoter. The value in *E. coli* wild-type was set to 1.

Statistical analysis: Error bars indicate SD; $n = 3$ biological replicates. Student's *t*-test was employed to assess the statistical significance of differences (*$p < 0.05$, **$p < 0.01$).

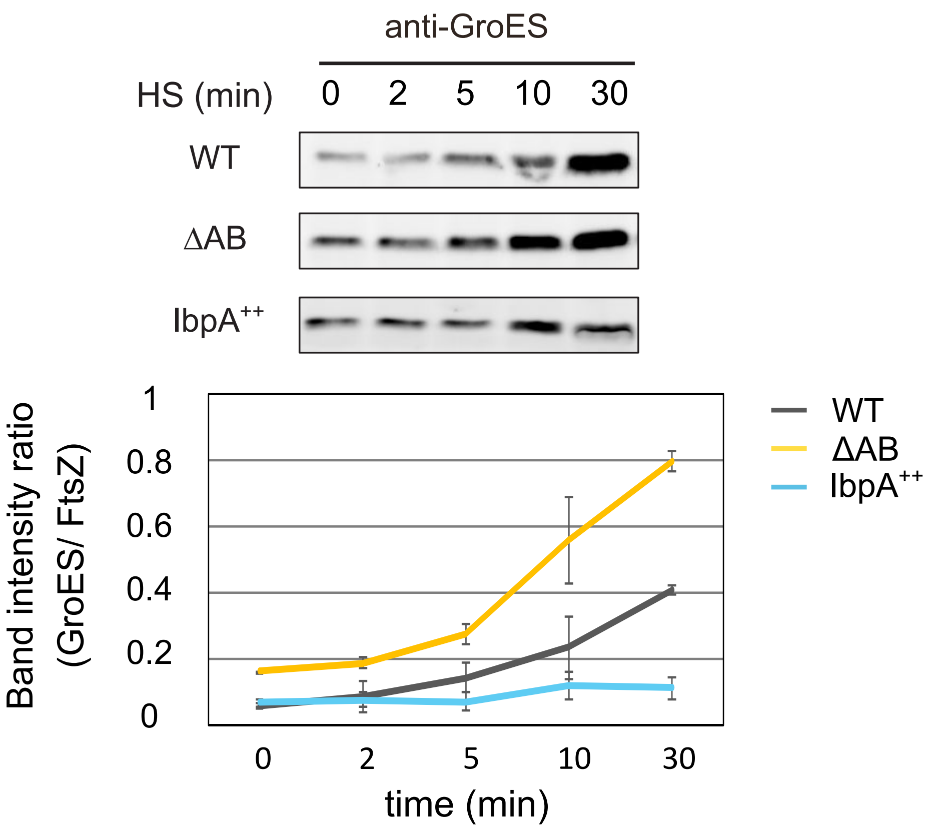


**Fig. S4. Time course of GroES levels following heat stress.**

*Top:* Western blotting of GroES in *E. coli* wild-type (WT), *ibpAB* operon-deleted cells (Δ*AB*), and *E. coli* Δ*AB* cells overexpressing IbpA (IbpA$^{++}$) after heat shock at 42 °C. Anti-GroES antibody was used for detection. *Bottom:* Quantification of GroES bands. The band intensities were normalized by FtsZ band intensities, which are shown in Fig. 5. HS (min); minutes from the start of heat shock. Error bars represent SD; *n* = 3 biological replicates.

**Table S1. *E. coli* strains used in this study**

| Strain | Genotype | Reference |
|---|---|---|
| DH5α | $F^-$, Φ80d*lacZ*ΔM15, Δ(*lacZYA-argF*) U169, *deoR*, *recA*1, *endA*1, *hsdR*17($r_K^-$,$m_K^+$), *phoA*, *supE*44, $\lambda^-$, *thi*-1, *gyrA*96, *relA*1 | Laboratory stock |
| BW25113 | Δ(*araD-araB*)567, Δ*lacZ*4787(::*rrnB*-3), $\lambda^-$ , *rph*-1, Δ(*rhaD-rhaB*)568, *hsdR514* | Laboratory stock |
| BL21 (DE3) | $F^-$ , *ompT*, *hsdS* ($rB^-$ $mB^-$), *gal*, *dcm* (λ DE3) | Laboratory stock |
| BW25113 Δ*ibpAB* | BW25113 Δ*ibpAB*::FRT-$Km^R$-FRT | (Miwa *et al*., 2021) |
| BW25113 Δ*dnaKJ* | BW25113 Δ*dnaKJ*::FRT-$Km^R$-FRT | (Miwa *et al*., 2021) |
| BL21 (DE3)Δ*ibpAB* | BL21 (DE3) Δ*ibpAB*::FRT-$Km^R$-FRT | (Miwa *et al*., 2021) |

**Table S2. Primers used in this study**

| Primer Name | | sequence |
|---|---|---|
| pBAD_ | vector_Fw | GGCCTATGCGGCCGCTAAGGG |
| pBAD_ | vector_Rv | ATGGAGAAACAGTAGAGAGTTGCGATAAAAAGCG |
| pBAD_ | sfGFP_Fw | agtaaaggagaagaacttttcactggag |
| pBAD_ | sfGFP_Rv | GCATAGGCCttatttgtatagctcatccatgcc |
| pBAD_ | sfGFP_vec_Rv | GCGGCCGCATAGGCCttatttgtatagctc |
| pBAD_ | dnaK_Fw | CTCTACTGTTTCTCCATACAACCACATGATGACCGAATATATAG |
| pBAD_ | dnaK_gfp_Rv | gaaaagttcttctcctttactacccaggtcgataccaattattttaccc |
| pBAD | dnaKpro_Fw | CCATAAGATTAGCGGATCCTACTTGATGACGTGGTTTACGACCCCATTTAG |
| dnaKpro | vec5UTR_Rv | ctgtgaatgtttattcaactgaTATTCGGTCATCATGTGGTTGTGAGTC |
| pBAD | groS_Fw | CTCTACTGTTTCTCCATACCAGCCGGGAAACCACGTAAG |
| pBAD | groS_gfp_Rv | gaaaagttcttctcctttactcacgcgatcatgcaatggac |
| pBAD_ | groESpro_Fw1 | GCCTCATCCCCATTTCTCTGGTCAtcagttgaataaacattcacagagacttttatgag |
| pBAD_ | groESpro_Fw2 | GATTAGCGGATCCTACTTGAAGGGGCGAAGCCTCATCCCCATTTCTCTGGTCA |
| pBAD_ | rpoH_Fw | TCTCTACTGTTTCTCCATgtagccgatgaggacgc |
| pBAD_ | rpoH-6xHis_Rv | gaaaagttcttctcctttactCATAATCAATAGCTCCT |
| pBAD | rpoH_Fw | gaataaacattcacagagacttttatgactgacaaaatgcaaagtttagctttagc |
| rpoH | I54N_Fw | ctaaaacgctgaatctgtctcacctgcggtttgttgttcatattgctcg |
| rpoH | I54N_Rv | gagacagattcagcgttttagctgcttccagatcgccatggtaatgcag |
| pCA_ | vector_Fw | GGGTCGACCTGCAGCCAAGCTTAATTAG |

| | | |
|---|---|---|
| pCA_ | vector_Rv | CATAGTTAATTTCTCCTCTTTAATGAATT |
| pCA_ | sfGFP_Fw | GGAGAAATTAACTATGagtaaaggagaagaac |
| pCA_ | sfGFP_Rv | GGCTGCAGGTCGACCCttatttgtatagctcatcc |
| pCA_ | ibpA_Fw | GAGGAGAAATTAACTatgCGTAACTTTGATTTATC |
| pCA_ | ibpA_Rv | GCTGCAGGTCGACCCttaGTTGATTTCGATACGGC |
| pCA_ | ibpA_IEI-AEA_Fw | GTgccGAAgccAACtaaGGGTCGACCTGCAGCCAAGC |
| pCA_ | ibpA_IEI-AEA_Rv | ttaGTTggcTTCggcACGGCGCGGTTTTTTCGCTTC |
| qRT | dnaK_Fw | tggtggtcagactcgtatgccaat |
| qRT | dnaK_Rv | attgctacagcttcgtccgggtta |
| qRT | groS_Fw | gcgtaaagaagttgaaactaaatctgc |
| qRT | groS_Rv | gcggcttcacttcgccattttcaag |
| qRT | rpoH_Fw | TCGTAATTATGCGGGCTATGG |
| qRT | rpoH_Rv | CAGTGAACGGCGAAGGAG |
| pCA | sbst_res_pBAD_Fw | CTGTCAGACCAAGTTTACTCATATATACTTTAGATTG |
| pCA | sbst_res_pBAD_Rv | CAATCTAAAGTATATATGAGTAAACTTGGTCTGACAG |
| PT0456 | | ATAAGGCTTGAAAAGTTCATTTCC |
| PT0457 | | AACGTGCCGAAATATCTTAAACAG |
| PT0071 | | AAATTGGGCAGTTGAAACCAGAC |
| PT0072 | | TACAGGTGCTCGCATATCTTCAACG |
| PM0195 | | gatgtttcgcttggtggtcgaatgggcagg |
| PM0196 | | cctgcccattcgaccaccaagcgaaacatc |

**Table S3. Plasmids used in this study**

| Plasmid | Feature | |
|---|---|---|
| pBAD30 | PBAD expression system, p15A, AmpR | (Guzman et al., 1995) |
| pCA24N | cloning vector, ColE1, CmR | (Kitagawa *et al.*, 2005) |
| pKJE7 | PBAD expression system carrying *dnaK-dnaJ-grpE*, pACYC Ori, CmR | Takara |
| pCA24N-sfGFP | pCA24N carrying super folder gfp | (Miwa *et al.*, 2021) |
| pCA24N-ibpA | pCA24N carrying *ibpA* | (Miwa *et al.*, 2021) |
| pCA24N-ibpA_AEA | pCA24N carrying *ibpA*_IEI::AEA | (Miwa *et al.*, 2021) |
| pBAD-dnaK'-sfGFP | pBAD30 carrying *dnaK* 5´UTR- super folder gfp | This study |
| pBAD-dnaKpro-sfGFP | pBAD30 carrying *dnaK* promoter- vector 5´UTR- super folder gfp | This study |
| pBAD-groES'-sfGFP | pBAD30 carrying *groES* 5´UTR- super folder gfp | This study |
| pBAD-groESpro-sfGFP | pBAD30 carrying *groES* promoter- vector 5´UTR- super folder gfp | This study |
| pBAD-rpoH-His | pBAD30 carrying *rpoH* 5´UTR- *rpoH*-His-Tag | This study |
| pBAD-rpoH_I54N-His | pBAD30 carrying *rpoH* 5´UTR- *rpoH* I54N- His-Tag | This study |
| pBAD-vec-rpoH'-His | pBAD30 carrying vector 5´UTR- *rpoH*-His-Tag | This study |
| pCA24N-Amp-ibpA | pCA24N carrying *ibpA,* CmR::AmpR | This study |
